# Identification of meteorological factors affecting migration of wild birds into miyazaki and its relation to circulation of highly pathogenic avian influenza virus

**DOI:** 10.1101/390922

**Authors:** Genki Arikawa, Maiku Abe, Mai Thi Ngan, Shuya Mitoma, Kosuke Notsu, Nguyen Thi Huyen, Eslam Elhanafy, Hala El Daous, Emmanuel Kabali, Junzo Norimine, Satoshi Sekiguchi

## Abstract

Aim of our study is to establish models for predicting the number of migratory wild birds based on the meteorological data. From 136 species of wild birds, which have been observed at Futatsudate in Miyazaki, Japan, from 2008 to 2016, we selected the potential high-risk species, which can introduce highly pathogenic avian influenza (HPAI) virus into Miyazaki; we defined them as “risky birds”. We then performed regression analysis to model the relationship between the number of risky birds and meteorological data. We selected 10 wild bird species as risky birds: Mallard (*Anas platyrhynchos*), Northern pintail (*Anas acuta*), Eurasian wigeon (*Anas penelope*), Eurasian teal (*Anas crecca*), Common pochard (*Aythya ferina*), Eurasian coot (*Fulica atra*), Northern shoveler (*Anas clypeata*), Common shelduck (*Tadorna tadorna*), Tufted duck (*Aythya fuligula*), and Herring gull (*Larus argentatus*). We succeeded in identifying five meteorological factors associated with their migration: station pressure, mean value of global solar radiation, minimum of daily maximum temperature, days with thundering, and days with daily hours of daylight under 0.1 h. We could establish some models for predicting the number of risky birds based only on the published meteorological data, without manual counting. Dynamics of migratory wild birds has relevance to the risk of HPAI outbreak, so our data could contribute to save the cost and time in strengthening preventive measures against the epidemics.

## Introduction

Highly pathogenic avian influenza (HPAI) virus causes avian influenza (AI), which is prevalent around the world. It was first reported from a goose farm in Guangdong Province of China, in 1996 [1–3]. Food and Agriculture Organization (FAO), World Organization for Animal Health (OIE) and World Health Organization (WHO) reported, mortality of over 150 million poultry and economic loss of billions of dollars, as results of HPAI outbreaks in Asia in 2003 and 2004 [4]. Human cases of AI infection have also been reported as a result of exposure to aerosol, large airborne droplets, and direct contact with infected birds [5–7]. HPAI virus causes a rapid onset of severe viral pneumonia and subsequently death with 60% mortality rate [7]. According to reported data, HPAI outbreaks engendered a severe economic impact on the poultry industry and public health [7,8].

Migratory wild birds are considered the natural hosts of AI virus [9,10]. Migratory wild birds infected with AI virus can transmit viral pathogens to other populations and subsequently to new areas [11]. Wild birds migrating from the Russian Far East, eastern Siberia, and eastern Mongolia, which were hotbeds of the virus, were considered to contribute to HPAI outbreaks in east Asia, southeast Asia, and southern Asia [11,12]. HPAI outbreaks have occurred seasonally in many countries. In Japan, HPAI outbreaks have occurred at intervals of several seasons (Table 1). Over 46% of HPAI outbreaks in Japan, from 2004 to 2016, happened in Miyazaki prefecture which is located in southern Japan between 30°21’39” and 32°50’20” N latitude and 130°42’12” and 131°53’09” E longitude (Fig 1a). HPAI outbreaks were related to the arrival of migratory wild birds, which are reservoirs of HPAI virus [13]. An increase in the number of migratory wild birds that arrive, causes an increase of HPAI outbreaks risk [14,15]. However, the relationship between arrival of migratory wild birds that are relevant to HPAI outbreaks, and meteorological factors has not been investigated. If meteorological factors are associated with the number of migratory wild birds, these factors can be used as predictive variables to employ preventive measures and efficient surveillance against HPAI. The aim of this study is to identify the predictive variables and use linear regression analysis to quantify the relationship between meteorological factors and migration of wild birds.

**Fig 1a.**
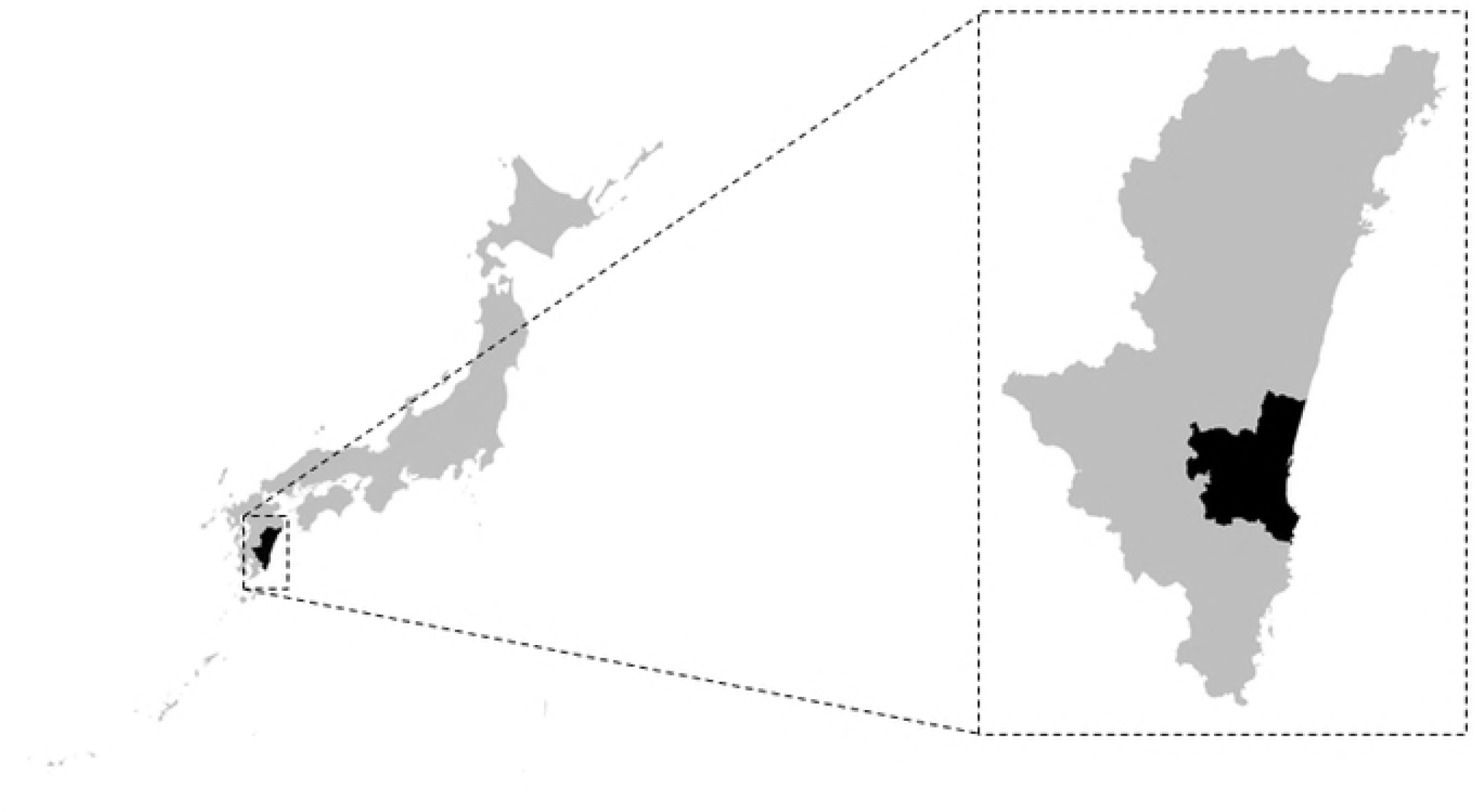
Location of Miyazaki prefecture. Miyazaki Prefecture is a prefecture of Japan located on the eastern coast of the island of Kyushu.

**Table 1.** HPAI outbreak cases for poultry in Japan (as of 31 March 2016)

| Year | Month | Province | Species | Cases | Destroyed | Virus type |
| --- | --- | --- | --- | --- | --- | --- |
| 2004 | January | Yamaguchi | Layer | 1 | 34640 | H5N1 |
| 2004 | February | Oita | Japanese bantam,<br>Domestic duck | 1 | 14 | H5N1 |
| 2004 | February~March | Kyoto | Layer | 2 | 240000 | H5N1 |
| 2007 | January | Miyazaki | Broiler breeder, Broiler | 2 | 70000 | H5N1 |
| 2007 | January | Okayama | Layer | 1 | 12000 | H5N1 |
| 2007 | February | Miyazaki | Layer | 1 | 93000 | H5N1 |
| 2010 | November | Shimane | Layer | 1 | 20000 | H5N1 |
| 2011 | January | Kagoshima | Layer | 1 | 8600 | H5N1 |
| 2011 | January | Aichi | Layer | 1 | 150000 | H5N1 |
| 2011 | January | Miyazaki | Broiler breeder | 1 | 10200 | H5N1 |
| 2011 | January | Miyazaki | Layer | 1 | 410000 | H5N1 |
| 2011 | January | Miyazaki | Broiler | 1 | 10000 | H5N1 |
| 2011 | January | Miyazaki | Broiler | 1 | 92000 | H5N1 |
| 2011 | January | Miyazaki | Broiler breeder | 1 | 6600 | H5N1 |
| 2011 | January | Miyazaki | Broiler | 1 | 40000 | H5N1 |
| 2011 | January | Miyazaki | Broiler | 1 | 40000 | H5N1 |
| 2011 | January | Miyazaki | Broiler | 1 | 190000 | H5N1 |
| 2011 | February | Aichi | Laying broiler breeder | 1 | 17500 | H5N1 |
| 2011 | February | Oita | Layer | 1 | 10000 | H5N1 |
| 2011 | February | Wakayama | Layer | 1 | 120000 | H5N1 |

**Table 1 HPAI outbreak cases for poultry in Japan (as of 31 March 2016) (continued)**
| Year | Month | Province | Species | Cases | Destroyed | Virus type |
| --- | --- | --- | --- | --- | --- | --- |
| 2011 | February | Mie | Broiler | 1 | 67000 | H5N1 |
| 2011 | February | Mie | Layer | 1 | 260000 | H5N1 |
| 2011 | February | Miyazaki | Broiler | 1 | 40000 | H5N1 |
| 2011 | February | Miyazaki | Broiler | 1 | 96000 | H5N1 |
| 2011 | February | Miyazaki | Broiler | 1 | 30000 | H5N1 |
| 2011 | February | Miyazaki | Broiler | 1 | 33000 | H5N1 |
| 2011 | February | Miyazaki | Broiler | 1 | 7500 | H5N1 |
| 2011 | February | Nara | Layer | 1 | 100000 | H5N1 |
| 2011 | March | Miyazaki | Broiler | 1 | 30000 | H5N1 |
| 2011 | March | Chiba | Layer | 1 | 35000 | H5N1 |
| 2011 | March | Chiba | Broiler | 1 | 62000 | H5N1 |
| 2014 | April | Kumamoto | Broiler | 1 | 110000 | H5N8 |
| 2014 | December | Miyazaki | Broiler | 1 | 4000 | H5N8 |
| 2014 | December | Miyazaki | Broiler | 1 | 42000 | H5N8 |
| 2014 | December | Yamaguchi | Broiler | 1 | 32000 | H5N8 |
| 2015 | January | Okayama | Layer | 1 | 200000 | H5N8 |
| 2015 | January | Saga | Broiler | 1 | 73000 | H5N8 |

## Materials and Methods

### Study design

We aimed to analyse the statistical relationships between the number of migratory wild birds and meteorological data. Based on the meteorological data from one month before the observation of migratory wild birds, we tried to find which specific meteorological factors affected their number. By understanding the relation to specific factors, we can use the meteorological data one month before their observation to predict their number efficiently (Fig 2).

**Fig 2.**
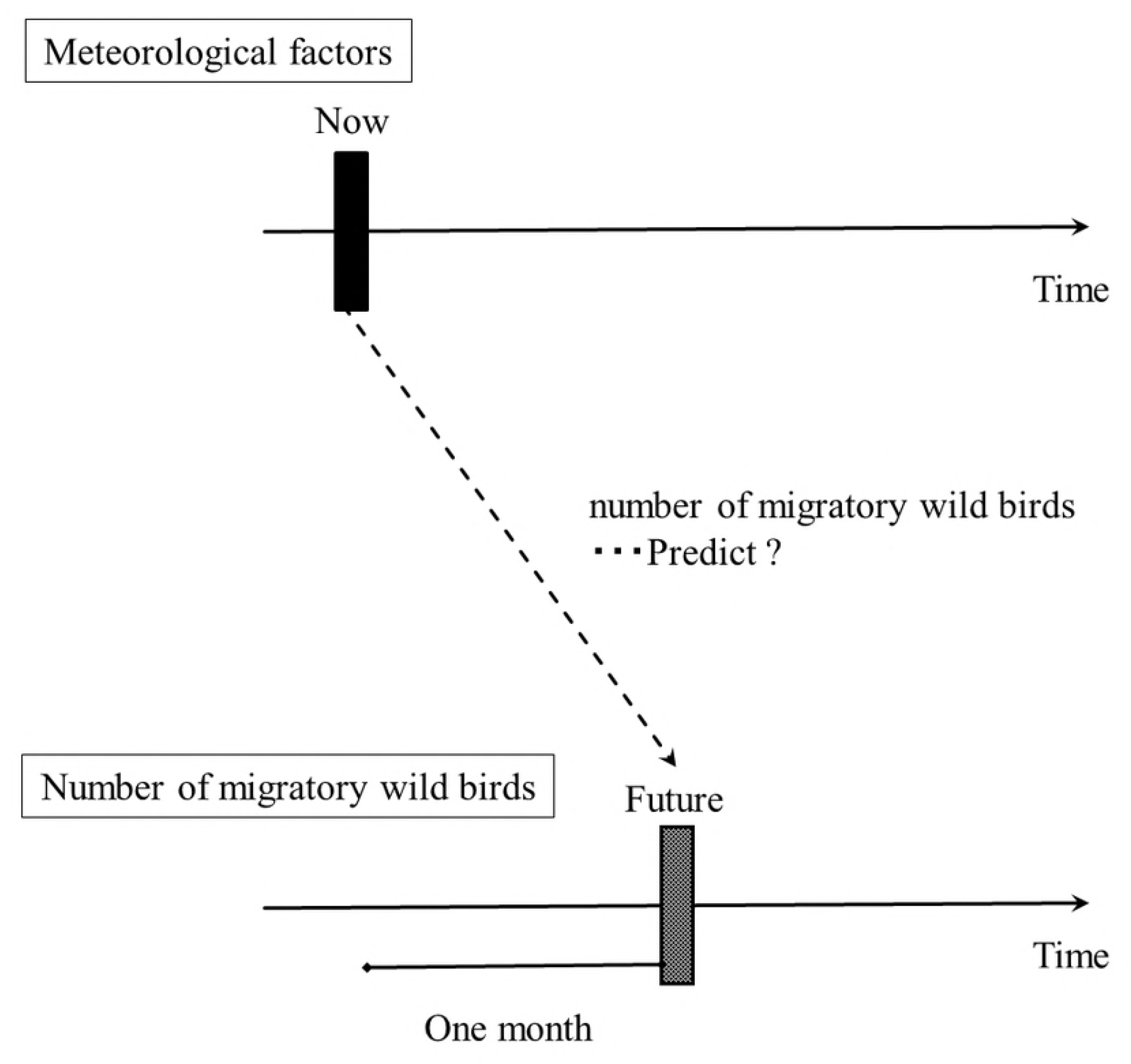
Study design. The upper figure shows the timeline for the meteorological factors. The lower figure shows the timeline for the number of wild birds. If we understand the relationship between meteorological factors and the number of wild birds, we can predict future number of wild birds from present meteorological data without manual counting.

### Data source for wild birds migrating into Miyazaki

The number of wild birds is recorded during migration period, from September to May every year, by the Ministry of Environment in Japan (http://www.env.go.jp/nature/dobutsu/bird_flu/migratory/ap_wr_transit/index.html). The purpose of this observation is to understand the tendency of wild bird species and the number of wild birds migrating to the wildlife sanctuary designated by the government during migration period. Recently, this data has also been used for employing HPAI outbreak preventive measures by the governments. This is open-source data and is updated monthly. In Japan, there are 39 observation points, which include two points in Miyazaki: Futatsudate and Miike. Futatsudate is located in the area, with most frequent HPAI outbreaks during the period spanning from January 2004 through March 2016. Most of HPAI outbreaks in Miyazaki have happened around Futatsudate (Fig 1b). Moreover, no HPAI outbreaks have happened around Miike during this period, so we decided to choose the data collected from Futatsudate point.

**Fig 1b.**
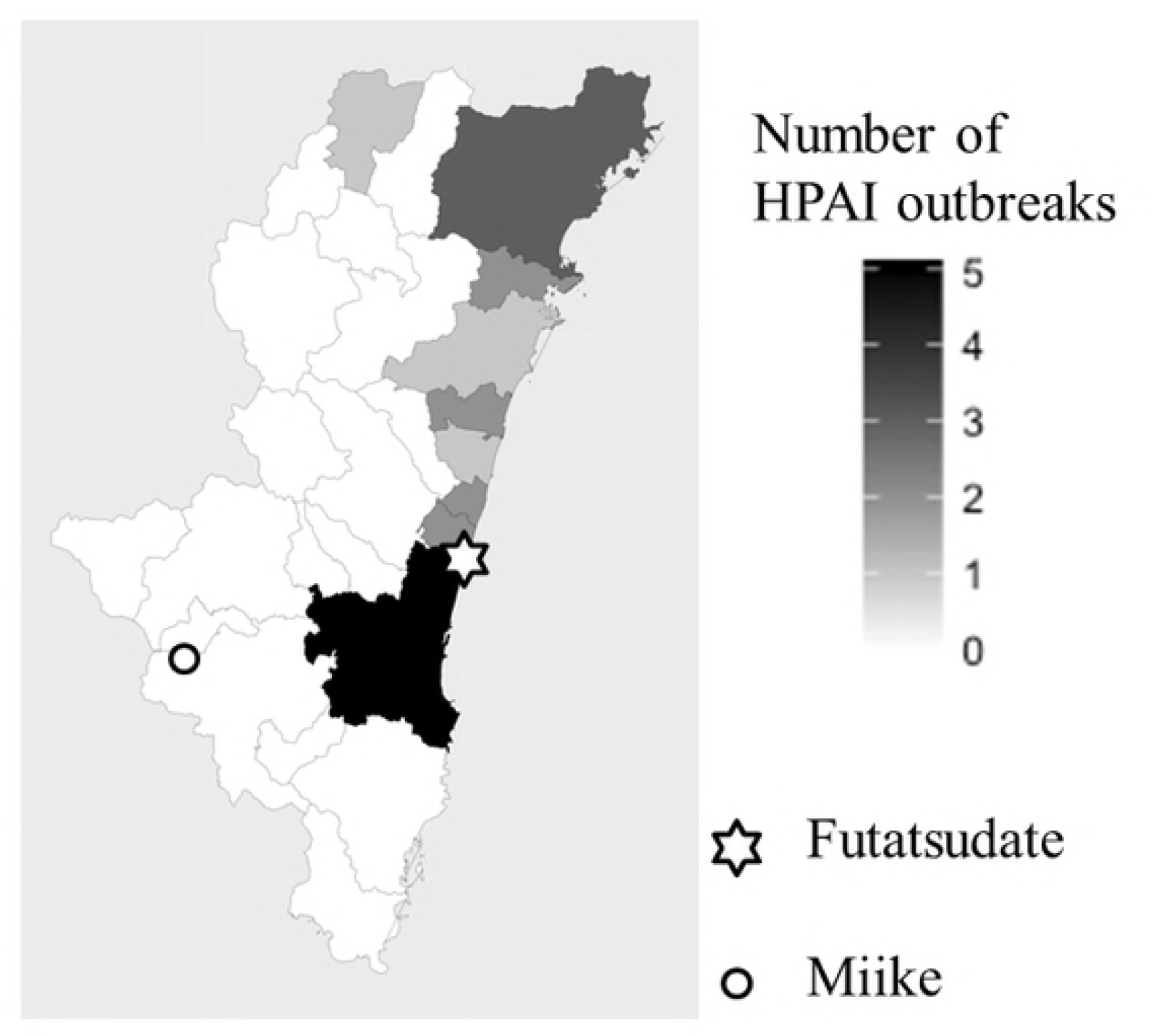
Location of Futatsudate and Miike. Futatsudate (white hexagram) is located in the center of the coastal area of Miyazaki City, at 32°03’37”N latitude and 131°49’53” E longitude. Miike (white circle) is situated in the west area of Miyazaki prefecture, at 31°72’35”5 latitude and 130°71’26” E longitude.

The data for 282 days for the period from June 2008 through March 2016, three observations per month, was used. Linear correction was used when there was a missing value.

### Selection of wild birds

The potential high-risk species were selected from all wild birds observed at Futatsudate during the research period. The selection of wild birds was based on the criteria reported by the European Safety Authority [16] (Fig 3). We followed the steps for selection of wild birds as described in Fig 3. In this study, we defined wild birds fitting in these criteria as “risky birds”. In the statistical analysis, total number of risky birds and individual number of risky birds of each species were used.

**Fig 3.**
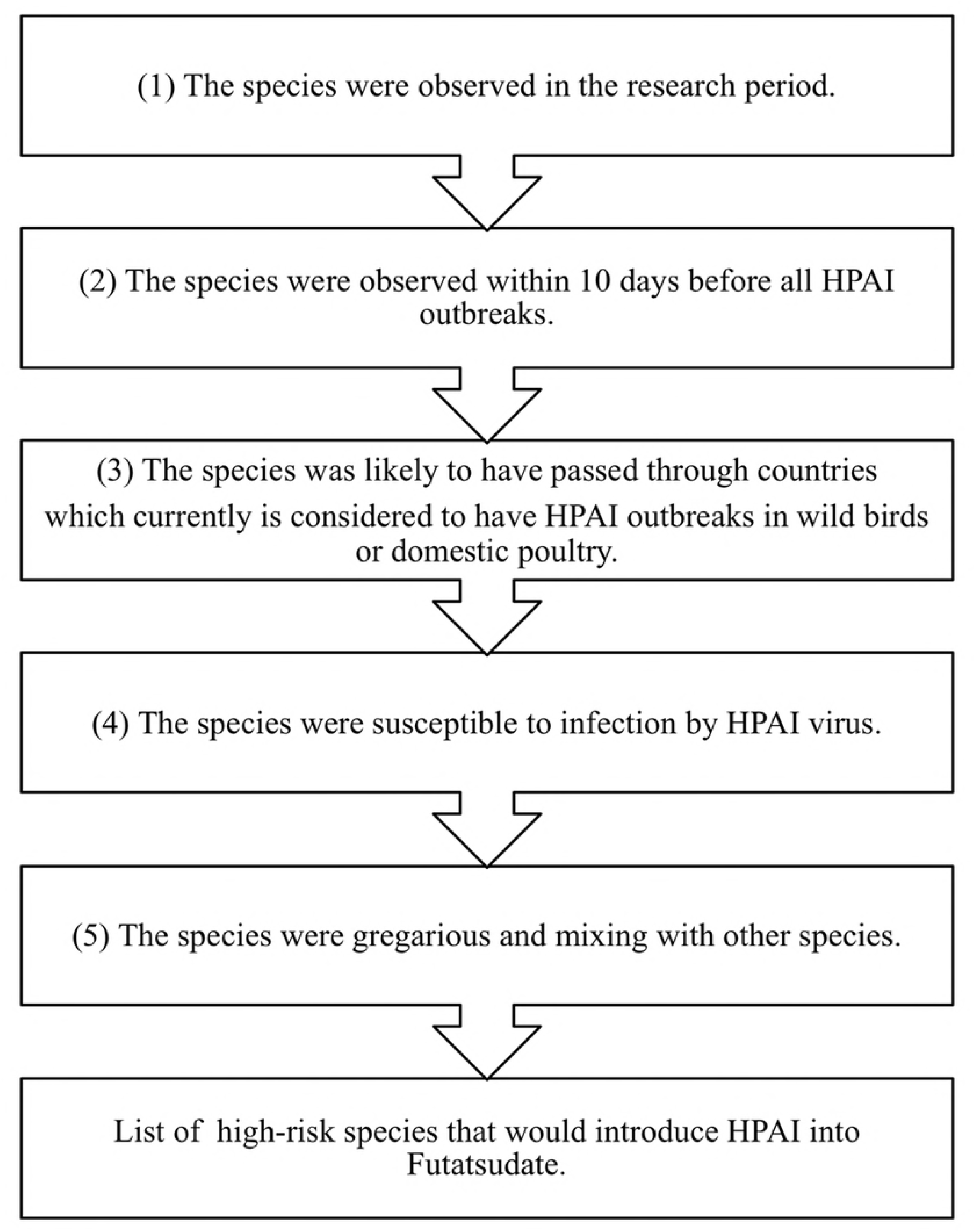
Decision tree for the selection of migratory wild birds more likely to introduce HPAI into Miyazaki. HPAI, highly pathogenic avian influenza. 136 species were checked for the first criteria. If any species meets first criteria, it will be passed to the next criteria. Finally, we considered the species met all criteria as a risky bird.

### Source of meteorological data

Open-source meteorological information is made available by the Japan Meteorological Agency (http://www.data.jma.go.jp/gmd/risk/obsdl/index.php). The meteorological factors such as atmospheric pressure, temperature, humidity, wind speed, precipitation, snow depth, sunshine hours, solar radiation, clouds, visibility, and atmospheric phenomena are observed by weather stations. Among these, precipitation, wind speed, temperature, and sunshine time are also observed by Automated Meteorological Data Acquisition System (AMeDAS). These data are updated every day. In Japan, there are about 60 weather stations and about 1,300 AMeDAS (http://www.jma.go.jp/jma/kishou/know/chijyou/surf.html; http://www.jma.go.jp/jma/kishou/know/amedas/kaisetsu.html). We used data, of about 10 days per month (early month, middle of month, and late month), for 36 factors of meteorological information, observed at the Miyazaki local weather station, from June 2008 through March 2016 (Table 2). This is the nearest local weather station to Futatsudate.

**Table 2.**
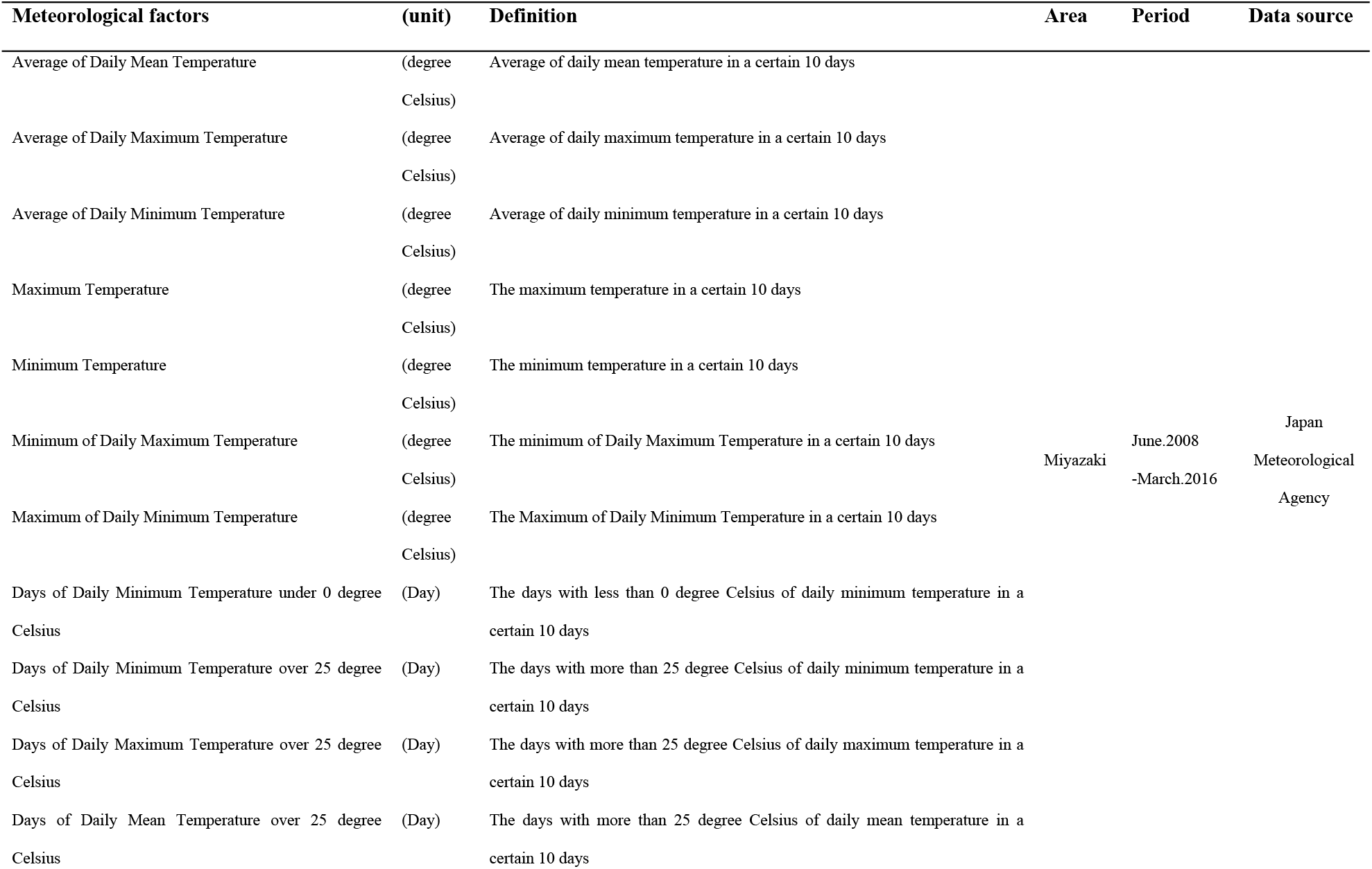
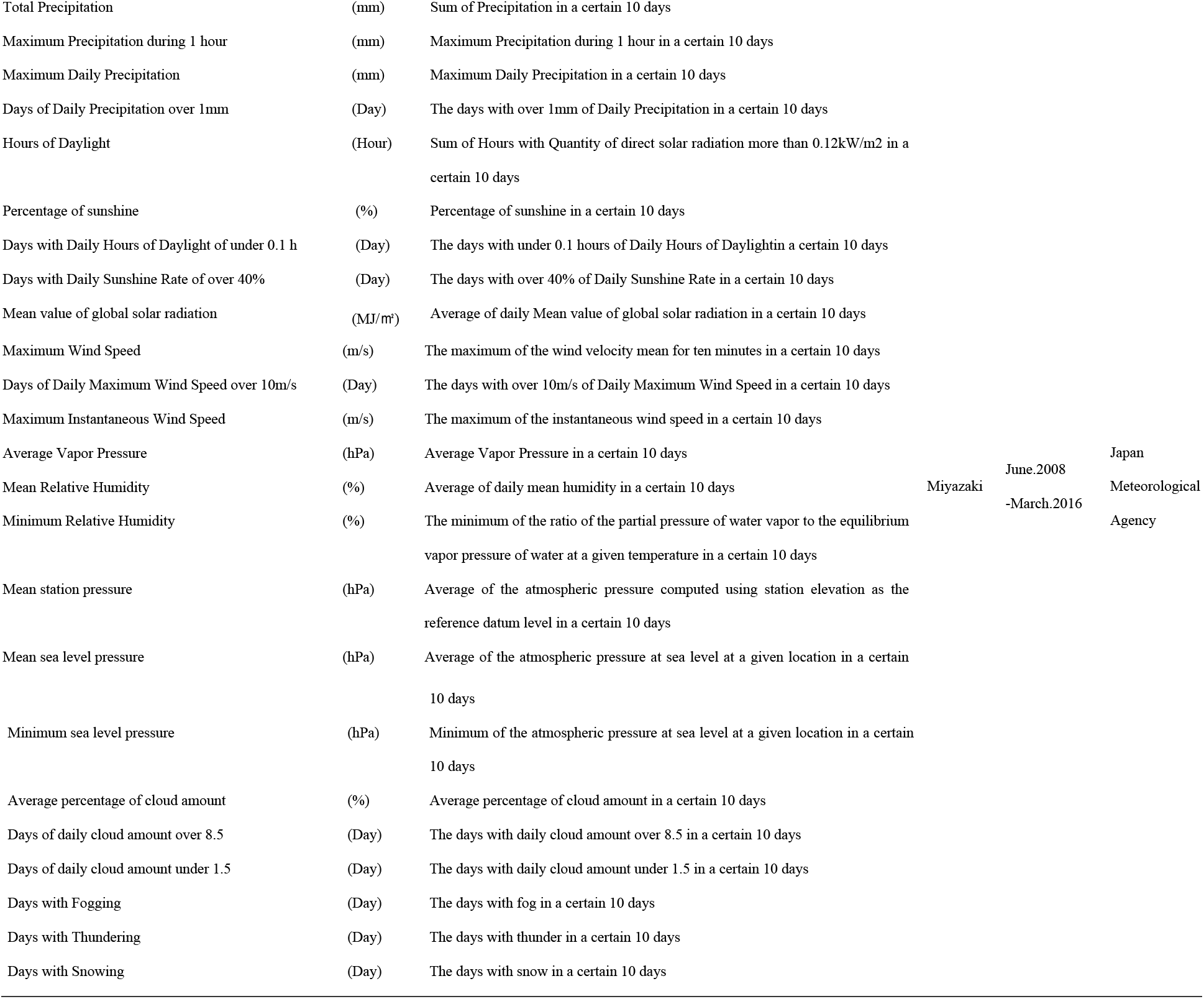
Define of Meteorological Factors.

| **Meteorological factors** | **(unit)** | **Definition** | **Area** | **Period** | **Data source** |
| --- | --- | --- | --- | --- | --- |
| Average of Daily Mean Temperature | (degree Celsius) | Average of daily mean temperature in a certain 10 days |  |  |  |
| Average of Daily Maximum Temperature | (degree Celsius) | Average of daily maximum temperature in a certain 10 days |  |  |  |
| Average of Daily Minimum Temperature | (degree Celsius) | Average of daily minimum temperature in a certain 10 days |  |  |  |
| Maximum Temperature | (degree Celsius) | The maximum temperature in a certain 10 days |  |  |  |
| Minimum Temperature | (degree Celsius) | The minimum temperature in a certain 10 days |  |  |  |
| Minimum of Daily Maximum Temperature | (degree Celsius) | The minimum of Daily Maximum Temperature in a certain 10 days | Miyazaki | June.2008<br>-March.2016 | Japan<br>Meteorological<br>Agency |
| Maximum of Daily Minimum Temperature | (degree Celsius) | The Maximum of Daily Minimum Temperature in a certain 10 days |  |  |  |
| Days of Daily Minimum Temperature under 0 degree Celsius | (Day) | The days with less than 0 degree Celsius of daily minimum temperature in a certain 10 days |  |  |  |
| Days of Daily Minimum Temperature over 25 degree Celsius | (Day) | The days with more than 25 degree Celsius of daily minimum temperature in a certain 10 days |  |  |  |
| Days of Daily Maximum Temperature over 25 degree Celsius | (Day) | The days with more than 25 degree Celsius of daily maximum temperature in a certain 10 days |  |  |  |
| Days of Daily Mean Temperature over 25 degree Celsius | (Day) | The days with more than 25 degree Celsius of daily mean temperature in a certain 10 days |  |  |  |

|  |  |  |
| --- | --- | --- |
| Total Precipitation | (mm) | Sum of Precipitation in a certain 10 days |
| Maximum Precipitation during 1 hour | (mm) | Maximum Precipitation during 1 hour in a certain 10 days |
| Maximum Daily Precipitation | (mm) | Maximum Daily Precipitation in a certain 10 days |
| Days of Daily Precipitation over 1mm | (Day) | The days with over 1mm of Daily Precipitation in a certain 10 days |
| Hours of Daylight | (Hour) | Sum of Hours with Quantity of direct solar radiation more than 0.12kW/m<sup>2</sup> in a certain 10 days |
| Percentage of sunshine | (%) | Percentage of sunshine in a certain 10 days |
| Days with Daily Hours of Daylight of under 0.1 h | (Day) | The days with under 0.1 hours of Daily Hours of Daylight in a certain 10 days |
| Days with Daily Sunshine Rate of over 40% | (Day) | The days with over 40% of Daily Sunshine Rate in a certain 10 days |
| Mean value of global solar radiation | (MJ/m<sup>2</sup>) | Average of daily Mean value of global solar radiation in a certain 10 days |

**Table 2 Define of Meteorological Factors (continued)**
| Meteorological factors | (unit) | Definition | Area | Period | Data source |
| --- | --- | --- | --- | --- | --- |
| Mean Wind Speed | (m/s) | Average of daily mean wind Speed in a certain 10 days |  |  |  |
| Maximum Wind Speed | (m/s) | The maximum of the wind velocity mean for ten minutes in a certain 10 days |  |  |  |
| Days of Daily Maximum Wind Speed over 10m/s | (Day) | The days with over 10m/s of Daily Maximum Wind Speed in a certain 10 days |  |  |  |
| Maximum Instantaneous Wind Speed | (m/s) | The maximum of the instantaneous wind speed in a certain 10 days |  |  |  |
| Average Vapor Pressure | (hPa) | Average Vapor Pressure in a certain 10 days |  |  | Japan |
| Mean Relative Humidity | (%) | Average of daily mean humidity in a certain 10 days | Miyazaki | June.2008 | Meteorological |
| Minimum Relative Humidity | (%) | The minimum of the ratio of the partial pressure of water vapor to the equilibrium vapor pressure of water at a given temperature in a certain 10 days |  | -March.2016 | Agency |
| Mean station pressure | (hPa) | Average of the atmospheric pressure computed using station elevation as the reference datum level in a certain 10 days |  |  |  |
| Mean sea level pressure | (hPa) | Average of the atmospheric pressure at sea level at a given location in a certain |  |  |  |

|  |  |  |
| --- | --- | --- |
|  |  | 10 days |
| Minimum sea level pressure | (hPa) | Minimum of the atmospheric pressure at sea level at a given location in a certain |
|  |  | 10 days |
| Average percentage of cloud amount | (%) | Average percentage of cloud amount in a certain 10 days |
| Days of daily cloud amount over 8.5 | (Day) | The days with daily cloud amount over 8.5 in a certain 10 days |
| Days of daily cloud amount under 1.5 | (Day) | The days with daily cloud amount under 1.5 in a certain 10 days |
| Days with Fogging | (Day) | The days with fog in a certain 10 days |
| Days with Thundering | (Day) | The days with thunder in a certain 10 days |
| Days with Snowing | (Day) | The days with snow in a certain 10 days |

### Statistical analysis

We analyzed the meteorological data one month before bird migration (previous month) and the number of risky birds (current month). A univariate linear regression analysis was performed to assess the relationship between meteorological factor and the number of risky birds (both the total of risky birds and the individual number of risky bird) seen one month after observing meteorological factors (*p* <0.25). Meteorological factors associated with the number of risky birds (both total and individual number of risky bird) were subsequently tested together in a multivariable linear regression analysis (*p* <0.05). A stepwise backward selection method was used to identify factors for final models. The fit of final models were assessed using Akaike information criterion. Multicollinearity was evaluated by variance inflation factor [17,18]. All statistical analyses were conducted by R software ver.3.2.1 (R development core team, Vienna, Austria).

## Results

### Selection of wild birds

Ten risky bird species were chosen from 136 species of wild birds observed at Futatsudate from June 2008 through March 2016 (see Appendix S1 in Supporting Information). Thirty-six wild bird species were observed whenever HPAI occurred in the research period and within 10 days before all HPAI outbreaks. Among these 36 species of wild birds, the species likely to have passed through countries considered to have HPAI outbreaks in wild birds or domestic poultry were 16. Among these, the species susceptible to HPAI viral infection were 12. There were two species of wild birds without any report about susceptibility to HPAI virus, but we decided to push forward those two species of wild birds to the next criterion, because of the possibility that these two species could be peculiar to Miyazaki and could bring HPAI virus into Miyazaki prefecture. In the final criterion, the number of wild bird species, which were highly gregarious and frequently mixing with other species were 10 (Table 3).

**Table 3.** List of 10species of risly birds used in this study.

| **Common name** | **Latine name** | **Order** |
| --- | --- | --- |
| Mallard | *Anas platyrhynchos* | Anseriformes |
| Northern pintail | *Anas acuta* | Anseriformes |
| Common pochard | *Aythya ferina* | Anseriformes |
| Eurasian wigeon | *Anas penelope* | Anseriformes |
| Eurasian teal | *Anas crecca* | Anseriformes |
| Eurasian coot | *Fulica atra* | Gruiformes |
| Common shelduck | *Tadorna tadorna* | Anseriformes |
| Northern shoveler | *Anas clypeata* | Anseriformes |
| Tufted duck | *Aythya fuligula* | Anseriformes |
| Herring gull | *Larus argentatus* | Charadriiformes |

### Descriptive statistics

The quantiles of the number of risky birds differed from each other (Table 4). These values for the 10 species were variable (for example, the mean value fluctuated from 7.7 to 192.2). During the research period, Northern pintail was the most frequently observed bird.

**Table 4.** Quartiles of number of 10 wild bird species used in this study.

| **Selected wild birds** | **Min.** | **1stQu.** | **Median** | **Mean** | **3rdQu.** | **Max.** |
| --- | --- | --- | --- | --- | --- | --- |
| All | 0.0 | 0.0 | 231.4 | 559.5 | 954.4 | 3288.0 |
| Mallard | 0.0 | 0.0 | 21.3 | 192.2 | 309.0 | 1457.0 |
| Northern pintail | 0.0 | 0.0 | 8.3 | 132.0 | 140.5 | 1985.0 |
| Eurasian wigeon | 0.0 | 0.0 | 20.8 | 63.3 | 65.0 | 911.0 |
| Eurasian teal | 0.0 | 0.0 | 24.0 | 56.0 | 77.0 | 325.0 |
| Common pochard | 0.0 | 0.0 | 11.4 | 49.7 | 83.0 | 340.0 |
| Eurasian coot | 0.0 | 0.0 | 13.5 | 20.2 | 34.0 | 94.0 |
| Northern shoveler | 0.0 | 0.0 | 6.0 | 15.4 | 26.5 | 83.0 |
| Common shelduck | 0.0 | 0.0 | 3.2 | 13.9 | 20.5 | 136.0 |
| Tufted duck | 0.0 | 0.0 | 3.2 | 9.4 | 15.8 | 55.0 |
| Herring gull | 0.0 | 0.0 | 2.5 | 7.7 | 8.8 | 118.0 |

Meteorological factors were classified into eight groups. Of the 36 meteorological factors, eleven were related to air temperature, four to precipitation, five to sunshine, four to wind speed, three to humidity, three to atmospheric pressure, three to clouds, and three were related to others (Table 5).

**Table 5.**
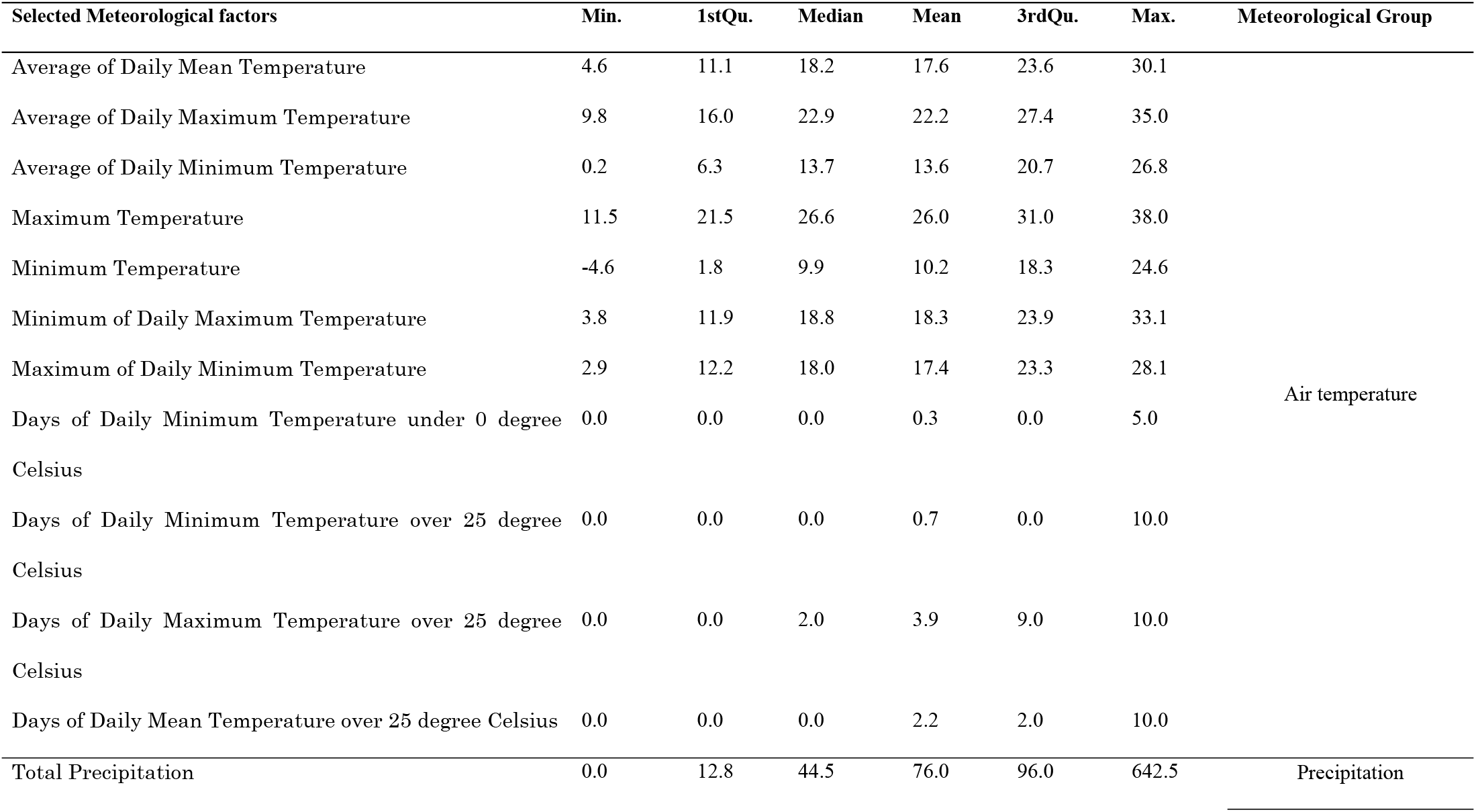
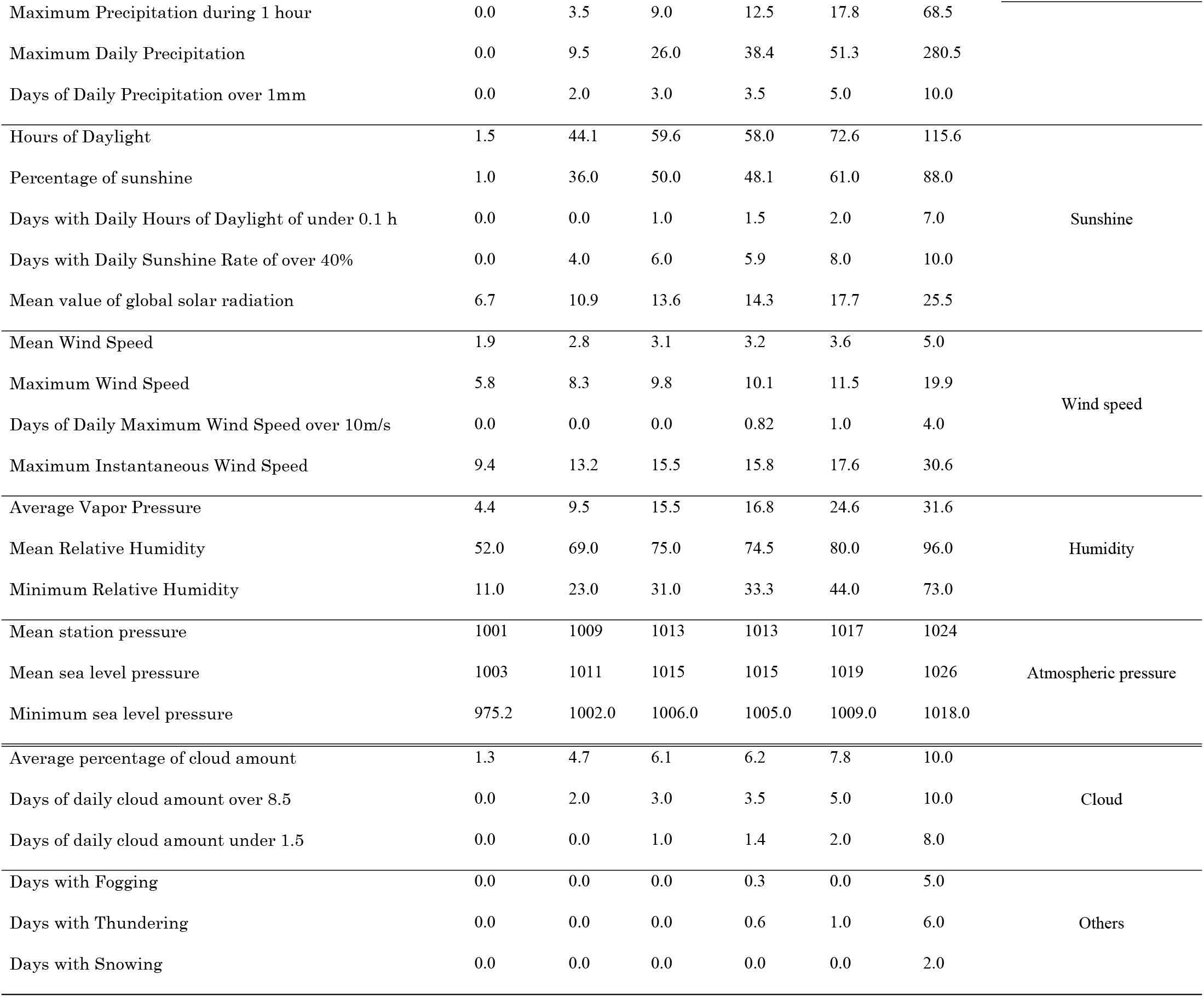
Quartiles of Meteorological Factors.

### Meteorological factors associated with the number of migrating risky birds

Many meteorological factors showed a significant association with the total number of risky birds (*p*<0.25) (Table 6) and the individual number of risky birds (*p*<0.25) (see Appendix S2). Air temperature variation, precipitation and humidity showed a significantly inverse association with the total number of risky birds. The group of factors related to sunshine showed both direct and inverse association with the total number of risky birds. Meteorological factors that affect the number of observations in the following month varied for each risky bird species. As results of multi-regression analysis, we could estimate the total number of risky birds in association with five meteorological factors (Table 7). To estimate the number of each risky bird, we needed one meteorological factor at least and five meteorological factors at most (see Appendix S3). Each risky bird needed a different meteorological factor to estimate the number, and there is no common meteorological factor to estimate individual number of risky birds of each species. Some meteorological factors were associated with the number of risky birds.

**Table 6.** A univariable association between Total Number of 10 selected wild bird species at Futatsudate and meteorological factors.

|  | **Estimate** | **Std. Error** | **t value** | **Adjusted R-squared** | **p-value** |
| --- | --- | --- | --- | --- | --- |
| Average of<br>Daily Mean<br>Temperature | -50.435 | 5.139 | -9.815 | 0.2553 | <0.0001 |
| Average of<br>Daily<br>Maximum<br>Temperature | -52.581 | 5.581 | -9.421 | 0.2399 | <0.0001 |
| Average of<br>Daily | -46.474 | 4.687 | -9.916 | 0.2593 | <0.0001 |
| Minimum |  |  |  |  |  |
| Temperature |  |  |  |  |  |
| Maximum | -58.608 | 5.879 | -9.969 | 0.2614 | <0.0001 |
| Temperature |  |  |  |  |  |
| Minimum | -40.486 | 4.229 | -9.573 | 0.2459 | <0.0001 |
| Temperature |  |  |  |  |  |
| Minimum of | -44.536 | 5.208 | -8.551 | 0.206 | <0.0001 |
| Daily |  |  |  |  |  |
| Maximum |  |  |  |  |  |
| Temperature |  |  |  |  |  |
| Maximum of | -53.124 | 5.447 | -9.753 | 0.2529 | <0.0001 |
| Daily |  |  |  |  |  |
| Minimum |  |  |  |  |  |
| Temperature |  |  |  |  |  |
| Days of Daily | 239.7 | 43.49 | 5.511 | 0.09556 | <0.0001 |
| Minimum |  |  |  |  |  |
| Temperature |  |  |  |  |  |
| under 0 |  |  |  |  |  |
| degree Celsius |  |  |  |  |  |
| Days of Daily | -121.29 | 22.9 | -5.295 | 0.08865 | <0.0001 |
| Minimum |  |  |  |  |  |
| Temperature |  |  |  |  |  |
| over 25 degree |  |  |  |  |  |
| Celsius |  |  |  |  |  |
| Days of Daily | -72.583 | 9.053 | -8.017 | 0.1854 | <0.0001 |
| Maximum |  |  |  |  |  |
| Temperature |  |  |  |  |  |
| over 25 degree |  |  |  |  |  |
| Celsius |  |  |  |  |  |
| Days of Daily | -70.64 | 10.06 | -7.024 | 0.1481 | <0.0001 |
| Mean |  |  |  |  |  |
| Temperature |  |  |  |  |  |
| over 25 degree |  |  |  |  |  |
| Celsius |  |  |  |  |  |
| Total | -2.0182 | 0.4424 | -4.562 | 0.06651 | <0.0001 |
| Precipitation |  |  |  |  |  |
| Maximum | -16.221 | 3.448 | -4.704 | 0.07064 | <0.0001 |
| Precipitation |  |  |  |  |  |
| during 1 hour |  |  |  |  |  |
| Maximum | -4.109 | 1.024 | -4.012 | 0.05149 | <0.0001 |
| Daily |  |  |  |  |  |
| Precipitation |  |  |  |  |  |
| Days of Daily | -109.85 | 17.67 | -6.217 | 0.1193 | <0.0001 |
| Precipitation |  |  |  |  |  |
| over 1mm |  |  |  |  |  |
| Hours of | 3.029 | 2.029 | 1.493 | 0.0044 | 0.1366 |
| Daylight |  |  |  |  |  |
| Percentage of | 13.286 | 2.279 | 5.831 | 0.1061 | <0.0001 |
| sunshine |  |  |  |  |  |
| Days with | -55.53 | 28.01 | -1.982 | 0.01043 | 0.04844 |
| Daily Hours of |  |  |  |  |  |
| Daylight of |  |  |  |  |  |
| under 0.1 h |  |  |  |  |  |
| Days with | 85 | 17.23 | 4.932 | 0.0774 | <0.0001 |
| Daily |  |  |  |  |  |
| Sunshine Rate |  |  |  |  |  |
| of over 40% |  |  |  |  |  |
| Mean value of | -79.477 | 8.596 | -9.246 | 0.2331 | <0.0001 |
| global solar |  |  |  |  |  |
| radiation |  |  |  |  |  |
| Mean Wind | 29.9 | 73.81 | 0.405 | -0.003016 | 0.6857 |
| Speed |  |  |  |  |  |
| Maximum | 9.047 | 19.065 | 0.475 | -0.002795 | 0.6355 |
| Wind Speed |  |  |  |  |  |
| Days of Daily | 47.15 | 40.07 | 1.176 | 0.00138 | 0.2404 |
| Maximum |  |  |  |  |  |
| Wind Speed |  |  |  |  |  |
| over 10m/s |  |  |  |  |  |
| Maximum | -6.968 | 11.941 | -0.584 | -0.002378 | 0.56 |
| Instantaneous |  |  |  |  |  |
| Wind Speed |  |  |  |  |  |
| Average | -42.56 | 4.62 | -9.213 | 0.2318 | <0.0001 |
| Vapor |  |  |  |  |  |
| Pressure |  |  |  |  |  |
| Mean Relative | -21.2 | 5.14 | -4.125 | 0.05447 | <0.0001 |
| Humidity |  |  |  |  |  |
| Minimum | -12.304 | 3.331 | -3.693 | 0.04349 | 0.0002667 |
| Relative |  |  |  |  |  |
| Humidity |  |  |  |  |  |
| Mean station | 79.539 | 6.763 | 11.76 | 0.3306 | <0.0001 |
| pressure |  |  |  |  |  |
| Mean sea | 78.604 | 6.711 | 11.71 | 0.3288 | <0.0001 |
| level pressure |  |  |  |  |  |
| Minimum sea | 39.782 | 6.028 | 6.600 | 0.1328 | <0.0001 |
| level pressure |  |  |  |  |  |
| Average | -168.82 | 20.22 | -8.347 | 0.1981 | <0.0001 |
| percentage of |  |  |  |  |  |
| cloud amount |  |  |  |  |  |
| Days of daily | -110.22 | 16.6 | -6.642 | 0.1343 | <0.0001 |
| cloud amount |  |  |  |  |  |
| over 8.5 |  |  |  |  |  |
| Days of daily | 199.11 | 25.47 | 7.817 | 0.1778 | <0.0001 |
| cloud amount |  |  |  |  |  |
| under 1.5 |  |  |  |  |  |
| Days with | -8.705 | 64.415 | -0.135 | -0.003544 | 0.8926 |
| Fogging |  |  |  |  |  |
| Days with | -246.27 | 38.91 | -6.329 | 0.1232 | <0.0001 |
| Thundering |  |  |  |  |  |
| Days with | 216.95 | 217.97 | 0.995 | -0.00003347 | 0.3204 |
| Snowing |  |  |  |  |  |

**Table 7.** Multivariable association between Total Number of 10 bird species at Futatsudate and meteorological factors hypothesized to influence that migrating into Miyazaki.

| Variable | Estimate | Std. Error | t value | p-value |
| --- | --- | --- | --- | --- |
| Mean station pressure | 49.008 | 10.362 | 4.73 | <0.0001 |
| Mean value of global solar radiation | -106.755 | 10.836 | -9.852 | <0.0001 |
| Minimum of Daily Maximum Temperature | 26.323 | 7.676 | 3.429 | 0.000698 |
| Days with Thundering | -143.639 | 34.528 | -4.16 | <0.0001 |
| Days with Daily Hours of Daylight of under 0.1 h | -172.246 | 26.88 | -6.408 | <0.0001 |
| **F-statistic** | 58.12 on 5 and 273 DF |  |  |  |
| **Adjusted R-squared** | 0.5067 | **p-value** | <0.0001 |  |

## Discussion and conclusions

The relationship between the number of migratory wild birds coming to Miyazaki and local meteorological data was assessed by linear regression analysis using open-source data of 282 days in Miyazaki, Japan. We selected 10 species of risky birds more likely to introduce HPAI virus into Miyazaki from 136 species. The migration of 10 species of risky birds into Miyazaki had a significant relation to five meteorological factors the month prior to their arrival. Our findings indicate that the number of migratory wild birds coming to Miyazaki in the next month could be predicted by using some meteorological data of Miyazaki in the present month. We think that predicting the number of migratory wild birds contributes to efficient preventive measure for the infectious diseases derived from migratory wild birds.

We succeeded in identifying five meteorological factors associated with migration of risky birds. These meteorological factors could be categorized into two main groups: air temperature and sunlight. First, there was a significantly inverse relationship between air temperature and the number of risky birds. Low temperature in autumn makes migratory wild birds migrate southwards to breed [19]. In a low temperature environment, it is difficult for birds to find food and water because plants cannot grow and water bodies could freeze. Additionally, severe weather conditions, especially low temperatures, are known to cause stress in birds [20–25]. Thus, in autumn, birds leave their breeding grounds, and migrate to a relatively warmer wintering ground at lower latitude to survive. Wintering grounds for the birds, including Japan, are located at a lower latitude than their breeding grounds, such as Siberia and Mongolia. Thus, migratory wild birds come to Japan in winter. This is why the number of migratory wild birds increases in Japan in winter. For the above reasons, air temperature, and the number of risky birds has a significantly inverse relationship in our study.

Second, there was a significantly direct relationship between the days with daily hours of daylight under 0.1 h and the number of risky birds. The climate of Miyazaki is finer, and less cloudy. Sunshine affects migration and distribution of wild birds [26–28], and plant growth [27]. Birds often gather in sunny areas such as Qinghai Lake [29] and search for areas with abundant plants. Miyazaki is also a famous prefecture with strong sunshine in Japan. We assert these facts as reasons for increase in the number of migratory wild birds in Miyazaki during winter.

Bird migration has an important role in the transmission and dissemination of several infectious diseases [11]. These diseases include HPAI, West Nile Virus, Lyme-Disease, and enteropathogens [11]. Particularly, HPAI is a threat to both animal and public health sectors [7,8].

Bird migration is, therefore, one of the important risk factors for HPAI outbreaks [30]. Actually, HPAI outbreaks have occurred in countries where many migratory wild birds migrate from breeding grounds. Japan is no exception. According to sequence data of HPAI virus, HPAI virus strains isolated in Japan and those that caused outbreaks in China and South Korea were genetically close [31,32]. The risk of HPAI outbreaks depends on the number of migratory wild birds [14,15,33]. Therefore, it is very important to understand the number of risky birds, which contributes to efficient preventive measures.

In Japan, every autumn, when migratory wild birds arrive, administrative organizations such as national and local governments strengthen measures to prevent HPAI outbreaks and spread of infectious diseases. Concrete preventive measures include (i) repairing the poultry houses so that wild birds and small animals cannot invade into the poultry houses, (ii) improve hygiene of the environment around poultry houses in order not to attract wild birds, (iii) keeping watch over people and vehicles entering and exiting the poultry houses, (iv) early detection and early reporting, (v) preparation of personnel and prevention materials in advance, and (vi) establishment of network between relevant organizations. Some local governments continue to strengthen preventive measures from October through April. However, mental burdens of local governmental staffs and poultry farmers are large. If it is possible to predict the number of migratory wild birds coming next month, administrative organizations will encourage poultry farmers to take strict biosecurity and appropriate preventive measures in advance. We could predict the number of migratory wild birds with our technique one month before migratory wild birds arrived. This makes it possible to switch from burdensome and ineffective preventive measures to smarter ones.

The meteorological factors may be different and species of migratory wild birds may vary greatly depending on the country and area. For example, in a country near the equator, temperatures are stable within high temperature range. As such, the number of migratory wild birds may not be affected by change of temperature. Observations of meteorological data and wild bird data are conducted in many countries. Our technique will be applicable to other areas in Japan and other countries. Our technique can predict the number of migratory wild birds using only the internet open-source data without watching and counting birds, which needs manpower and expertise (e.g., how to distinguish bird species and how to count groups of birds). In this study, weather factors influencing migration of wild birds were identified. In the future study, we would like to conduct spatial analysis using meteorological factors in the survey area, the number of poultry farms, the distance to reservoir, and so on.

In conclusion, our technique enabled us to predict the number of migratory wild birds without counting. As a result, we could foretell the number of migratory wild birds coming to Miyazaki using open-source local meteorological data on the Internet. We suggest that this approach can be applied all over the world to predict the periods with high risk of HPAI outbreak in specific areas.

## Acknowledgements

The authors received no directed funds to conduct this research. We thank Naoaki Misawa, Yoshitaka Goto, Nariaki Nonaka, Tamaki Okabayashi, and Shinji Watanabe for comments and suggestions that helped improve the research.

We would like to thank Editage (www.editage.jp) for English language editing.

## Supporting information

**S1 Table. 136 species of wild birds which were observed from 2008 to 2016.**

**S2.1 Table. A univariable association between Number of Mallard at Futatsudate and meteorological factors.**

**S2.2 Table. A univariable association between Number of Northern pintail at Futatsudate and meteorological factors.**

**S2.3 Table. A univariable association between Number of Eurasian wigeon at Futatsudate and meteorological factors.**

**S2.4 Table. A univariable association between Number of Eurasian teal at Futatsudate and meteorological factors.**

**S2.5 Table. A univariable association between Number of Common pochard at Futatsudate and meteorological factors.**

**S2.6 Table. A univariable association between Number of Eurasian coot at Futatsudate and meteorological factors.**

**S2.7 Table. A univariable association between Number of Northern shoveler at Futatsudate and meteorological factors.**

**S2.8 Table. A univariable association between Number of Common shelduck at Futatsudate and meteorological factors.**

**S2.9 Table. A univariable association between number of Tufted duck at Futatsudate and meteorological factors.**

**S2.10 Table. A univariable association between number of Herring gull at Futatsudate and meteorological factors.**

**S3 Table. Multivariable association between number of each risky birds at Futatsudate and meteorological factors hypothesized to influence the migrating into Futatsudate.**

